# The role of vibration amplitude in the escape hatching response of red-eyed treefrog embryos

**DOI:** 10.1101/2024.09.23.614602

**Authors:** Julie Jung, Michael S. Caldwell, J. Gregory McDaniel, Karen M. Warkentin

## Abstract

The function and adaptive significance of defensive behaviors depend on the contexts in which they naturally occur. Amplitude properties of predator cues are widely used by prey to assess predation risk, yet rarely studied in the context of the stimuli relevant to defensive decisions in nature. Red-eyed treefrog embryos, *Agalychnis callidryas*, hatch precociously in response to attacks on their arboreal egg clutches by snakes and wasps. They use vibrations excited during attacks to detect predators, but wind and rainstorms also excite intense vibrations. Past work has demonstrated that to avoid costly decision errors, *A. callidryas* non-redundantly combine information from the temporal and frequency properties of clutch vibrations. Here we demonstrate that embryos also use absolute amplitude and fine-scale amplitude modulation information to refine their hatching decision. We used vibration recordings to characterize the amplitude properties of the most common predator and benign-source disturbances to *A. callidryas* egg clutches in nature and tested whether embryos at three ages across the onset of mechanosensory-cued hatching (4–6 days) respond to amplitude variation during playback of synthetic vibrations to eggs. Older embryos responded to much lower-amplitude vibrations, reflecting a >88-fold decrease in response threshold from 4 to 5 days. To assess how embryos combine amplitude with other vibration properties, we played embryos recorded exemplars of snake attack and rain vibrations of varying amplitudes and patterns of amplitude modulation. The amplitude response curve was steeper for snake recordings than for rain. While amplitude information alone is insufficient to discriminate predator attack from benign-source vibrations, *A. callidryas* employ an impressively complex strategy combining absolute amplitude, amplitude modulation, temporal, and frequency information for their hatching decision.

## Introduction

Stimulus amplitude is one of the major features that animals use to extract information from acoustic signals and cues, along with frequency, temporal pattern, and phase. The importance of amplitude variation in communication signals is well established (Gerhardt and Huber, 2002). Many animals – from natterjack toads (Arak, 1983) and gray treefrogs (Fellers, 1979a; Fellers, 1979b) to mole crickets (Forrest, 1980; Forrest, 1983) and eastern towhees (Nelson, 2000) – use amplitude modulation to encode information within their signals or extract information about signaler distance from the attenuation of signals (Searcy and Andersson, 1986). We know less about the roles the amplitude characteristics of incidental predator cues play in defensive decisions, and even less about how those defensive responses change with development.

Best studied is the use of cue amplitude information in the defensive startle responses displayed by a wide range of animals following exposure to brief, intense sound or vibrational stimuli (Eaton, 1984; Friedel, 1999). Most of this work, however, has used the startle response as a convenient tool with which to study other aspects of cognition, such as habituation, neural architecture, and learning (Eaton, 1984). While there are several notable exceptions in insect models (Faure and Hoy, 2000; Fullard et al., 2008; Hergenröder and Barth, 1983; Miller and Surlykke, 2001), for many of these studies, amplitude-dependent variation in defensive behavior is largely removed from contexts relevant to the evolution of decision strategies for antipredator defenses. By contrast, a related predator cue property, the intensity, or concentration, of chemical cues associated with predation risk has been studied extensively, with particular focus on the variation of cue intensity in the natural environment of prey, and how chemical cue intensity influences defensive decisions (Kats and Dill, 1998; Kusch et al., 2004; Schoeppner and Relyea, 2008; van Buskirk and Arioli, 2002).

Much as the intensity of chemical cues is often proportional to local predation risk, the amplitude of sound and substrate vibration cues might be expected to correlate well with the size, proximity, and activity level of predators. For small arthropods at least, the amplitude of incidental sounds produced while moving over leaf litter increases with animal size and speed of movement (Goerlitz et al., 2008), and such sound can be expected to attenuate reliably over distance (Bradbury and Vehrencamp, 1998). Prey would benefit if they were able to extract this sort of information from predator cues.

Indeed, studies generally show lower latency or stronger startle responses to stimuli of greater amplitude (Eaton, 1984), which could be indicative of larger or more immediate threats. The wandering spider, *Cupiennius salei* shows either predatory or defensive behavior depending on the amplitude properties of vibrational disturbances transmitted through plant substrate (Hergenröder and Barth, 1983). Still, it is not well understood how variation in prey response to stimuli of varying amplitudes is correlated with either actual predation risk or the overall amplitude or amplitude modulation characteristics of natural predator cues. Moreover, it is still not well understood how variation in prey response to stimuli of varying amplitudes is correlated with ontogenetic change, which may be due to sensory development (increasing sensitivity to lower amplitudes), changing decision rules (less costly false alarms) or both.

Using a combination of vibration recordings and playback experiments, we characterized the amplitude properties of vibrational disturbances to red-eyed treefrog egg clutches and tested the defensive response of embryos to variation in the amplitude characteristics of vibrational disturbance. Red-eyed treefrogs, *Agalychnis callidryas*, lay their eggs on vegetation overhanging ponds in the lowland wet forests of Central America. Although these eggs normally take 6–7 days to hatch, if attacked by an egg predator, such as a snake or wasp, they can hatch up to 30% precociously, escaping to the pond below (Warkentin, 1995). This defensive escape-hatching is cued by vibrations in the egg clutch excited by the attack (Warkentin, 2005), perceived by inner ear mechanoreceptors (Jung et al., 2019) and lateral line neuromasts (Jung et al., 2020), which play key roles in mediating the hatching response. Both motion-only and egg-contact cues stimulate the vestibular system, while contact cues additionally stimulate the lateral line neuromasts (Jung et al., 2020). Harmless wind and rain disturbances also excite intense vibrations within the clutch, with frequency distributions that largely overlap those excited by predators (Caldwell et al., 2009; Warkentin, 2005), and early hatchlings suffer increased mortality from aquatic predators (Warkentin, 1995; Warkentin, 1999; Willink et al., 2014). Embryos combine information from at least three temporal and two frequency characteristics of clutch vibrations to refine their hatching response, thus reducing the incidence of false alarms during benign disturbances (Caldwell et al., 2009; Caldwell et al., 2010; Jung et al., 2021; Warkentin et al., 2006b).

To determine how *A. callidryas* incorporate amplitude information into their hatching decision, we first categorized the amplitude properties of the most common vibrational disturbances to egg clutches at our field site in Soberania National Park, Panama, including attacks by four species of snake predators, one species of egg-eating wasp, wind and rain disturbances, and the vibrations generated by routine movements of embryos within their eggs. An earlier study which examined a smaller subset of these disturbance types found that egg-clutch vibrations excited by hard rain can be of higher amplitude than those excited during attacks by two species of snake (Warkentin, 2005). Next, we used playback of synthetic vibrations to examine amplitude-dependent variation in embryo hatching response and to study how this changes over the period of mechanosensory-cued hatching period (4–6 days). To assess if the effect of amplitude on the escape-hatching response varies with other cue properties, and how amplitude variation in natural egg-clutch disturbances affects escape hatching, we conducted playback experiments varying the amplitude of several recorded rainstorm and snake attack exemplars. Finally, to test whether *A. callidryas* responds to fine-scale amplitude modulation of disturbance cues, we played embryos stimuli that varied in amplitude envelope shape. We hypothesize that embryos rely on amplitude information to refine their hatching decisions and that development improves detection of lower amplitude stimuli. This research elucidates the roles of perceived amplitude information and sensory development and how they are related in the context of a critical defense behavior.

## Methods

### Egg collection and care

Experiments in this study were conducted between 2004 and 2017. In 2004 and 2005, we collected recently laid *A. callidryas* egg clutches and the leaves on which they were laid from Ocelot Pond, ∼2 km southeast of Gamboa, Panama (9.120894 N, 79.704015 W). In 2016 and 2017, we collected clutches from the Experimental Pond in Gamboa, located adjacent to the laboratory facilities at the Smithsonian Tropical Research Institute. Since most clutches are laid between 10 pm and 2 am (Warkentin, 2002; Warkentin et al., 2005), we assigned embryos to daily age-classes and report developmental timing of individuals starting from midnight of their oviposition night.

Clutches 0–3 days post oviposition were brought to an open-air ambient temperature and humidity laboratory in Gamboa. We removed any dead eggs or debris from the clutches, mounted them on plastic cards for support, and suspended them over cups of aged tap water in plastic cups. Clutches were misted with rainwater frequently to maintain hydration. All embryos used in experiments were morphologically normal, in developmental synchrony with siblings in their clutch, and in intact, turgid eggs at the start of the experiment. In a few experiments, some hatchlings (N = 18 individuals from 2 clutches) were preserved for morphological studies, to be presented elsewhere; all other hatchlings were returned to their respective ponds. This research was conducted under permits from the Panamanian Environmental Ministry (ANAM: SE/A-50-04, SE/A-49-05; MiAmbiente: SE/A-59-16, SE/A-55-17) and approved by the Institutional Animal Care and Use Committees of Boston University (02-013, 05-022, and 14-008) and the Smithsonian Tropical Research Institute (2014-0601-2017, 2017-0601-2020).

### Amplitude characteristics of common natural vibrational disturbances to egg clutches

The recording of substrate vibrations excited by predator and benign-source disturbances followed methods previously employed for *A. callidryas* egg clutches, and frequency characteristics of some of the recordings were analyzed elsewhere (Caldwell et al., 2009). Vibrations excited during rainstorms and predator attacks result from direct physical disturbances to the clutch. For most egg clutches, to improve ease of handling, reduce background vibrations from outside the clutch, and facilitate recording, we taped their leaves to plastic cards, and taped these cards to bricks or large jars filled with water. Because the strongest vibrations excited by wind result from forcing and movements of the plant substrate, we left all clutches used for wind recording attached to ∼50 cm of the plant on which they were laid. These plant sections were then rigidly attached to a heavy jar filled with water. We also left several rain and predator recording clutches attached to sections of their original oviposition substrate. For some types of direct forcing applied to the clutch, very low frequency vibrations (<10 Hz) differed between clutches mounted on jars and those attached to plants, while higher frequency vibrations did not appear to be affected. However, wind vibration recordings from clutches on ∼50 cm of plant substrate were similar to wind recorded from clutches in their original oviposition sites on intact plants (Caldwell, 2010).

To record natural vibrations from substrates, we embedded a small accelerometer [Endevco 25B (0.2 g), San Juan Capistrano, CA, USA or AP Technology AP19 (0.14 g), Oosterhout, Netherlands] within the jelly of each test clutch prior to hatching competence. Recordings were made with hatching competent, 4- to 5-day-old clutches. Accelerometers added ∼5% to the mass of each clutch, and test clutches remained within the natural range of inter-clutch mass variation (Warkentin, 2005). Transduced vibrational signals were amplified using a signal conditioner (Endevco 4416B or AP Technology APC7), digitized with an external sound card (MSE-U33HB, Onkyo USA, Saddle River, USA), and output was recorded using Canary bioacoustics software (v.1.2.4, Cornell University Laboratory of Ornithology, Ithaca, USA) at 22.1 kHz on a Macintosh G4 notebook computer. We calibrated the output of our recording setup using a representative accelerometer and a sinusoidal vibration stimulus of known amplitude. We then calculated the amplitude of each recording using the sensitivity of each individual accelerometer, as measured at the factory.

To record snake attacks, we collected snakes [*Leptophis ahaetulla*, *Leptodeira rhombifera*, (formerly *L. annulata*), *Leptodeira ornata* (formerly *L. septentrionalis*, (Barrio-Amorós, 2019; Torres-Carvajal et al., 2020) and *Imantodes inornatus*] from ponds near Gamboa. Snakes were housed in mesh cages under ambient temperature and humidity, and were provided vegetation, water, and *A. callidryas* egg clutches as food. Once snakes adjusted to captivity and began to feed regularly, they were given clutches with embedded accelerometers to record attack vibrations. Attacks were filmed using an infrared capable camera (Sony TRV120 or TRV250, New York, USA). After several successful recordings, snakes were released at their respective collection sites.

To record wasp attack vibrations, we trained wasps [*Polybia rejecta*] to return to a feeding station (Warkentin et al., 2006a). We marked wasps with dots of paint for individual identification. They were presented with *A. callidryas* egg clutches containing embedded accelerometers, and we recorded video and vibrations of attacks.

To record clutch vibrations excited by rain, we placed clutches containing accelerometers outdoors, in the path of unobstructed falling rain. While rain recordings from plant-mounted clutches may have included low-amplitude vibration produced by wind, we did not include recordings with periods of strong wind in our analysis.

To record wind, we placed plant sections with clutches containing embedded accelerometers outside during periods of particularly strong wind gusts, but without rain, and recorded clutch vibrations.

### Analysis of common natural vibrational disturbances to egg clutches

For predator attacks, we synchronized video and vibration recordings so that we could identify vibrations excited by specific predator and embryo behaviors. We then sampled vibrations associated with these events for analysis. For rain and wind recordings, video was not necessary to identify the source of vibrations. All measurements were made in Canary (v.1.2.4, Cornell University Laboratory of Ornithology, Ithaca, NY, USA). To quantify the overall intensity of predator attacks, rain, and wind recordings, we measured peak and root-mean-square (RMS) amplitudes from samples including long periods of intermittent vibrations recorded during each disturbance type. These samples included vibration excited by routine embryo movements and hatching as well as periods of vibrational silence that were recorded during the disturbances.

For snake attacks, samples began when the snake first attacked the egg clutch and ended following the last bout of embryo hatching not succeeded by additional hatching within 30 seconds of by any subsequent attack vibrations. We edited out any periods of vibrational silence lasting more than two minutes. During many attacks, snakes bit and attempted to swallow embedded accelerometers. We edited out any periods of accelerometer biting from the samples, and stopped sampling if the accelerometer was pulled from the clutch. Snake samples were 172 ± 142 seconds in length (mean ± s.d.).

Sample sizes per species were as follows: *L. ahaetulla*, seven individual snakes, 17 egg clutches attacked, 1–5 clutches per snake; *L.rhombifera* , seven snakes, 11 clutches, 1–4 clutches per snake; *L. ornata*, five snakes, 13 clutches, 1–4 clutches per snake; and *I. inornatus*, two snakes, one clutch each. A different analysis of recordings from two of the *L. ahaetulla* and three of the *L. rhombifera* used here was presented previously (Warkentin, 2005). All snake attacks sampled elicited escape hatching.

When sampling wasp attack vibrations, we considered an attack to begin when a wasp started biting at or pulling on the eggs, and to end following the last bout of embryo hatching not succeeded by additional hatching within 30 seconds or by any subsequent attack vibrations within two minutes. Wasps often make many short visits to a clutch, attacking it repeatedly over the course several hours until all of the embryos have hatched or have been consumed (Warkentin, 2000). Therefore, to obtain sufficiently long samples of wasp attack vibrations, we combined recordings of several attacks on each clutch, often by more than one wasp. For each clutch used in our analysis, we sampled between 47 and 300 seconds of attack vibrations depending on availability (261 ± 71 seconds, N = 18 clutches, 13 individual wasps, 1–3 periods of attack per wasp).

While low intensity wind and rain do occur in Gamboa (e.g. light breeze and drizzle), higher intensity benign disturbances are likely more difficult to discriminate from predator attacks. We therefore did not sample very low amplitude wind or rain vibrations. All benign disturbances included were, however, well within the intensity range commonly experienced by clutches observed at our field sites in Soberania and Gamboa, Panama. Rain samples from each clutch include the 300 s of vibration with the highest RMS amplitude (N = 19 clutches, 19 rainstorms). From each wind recording we also sampled the 300 seconds with the highest RMS amplitude (N = 7). Vibrations excited by routine movements of embryos within their eggs are brief and irregularly spaced. It was therefore not possible to accurately quantify amplitude of these common disturbances over timescales similar to those of the long-period predator, rain, and wind samples.

It is not known over what timescale embryos assess the amplitude properties of clutch vibrations. In addition to the longer samples of intermittent vibrational disturbance, we also measured the RMS amplitude of shorter periods of continuous vibration (RMScont), without the periods of vibrational silence that occur during most disturbances. For these shorter samples, we began with the highest RMS sections of each longer sample described above, and edited out any sections with amplitudes below the noise floor of our equipment (0.10 m/s^2^peak, 0.026 m/s^2^ RMS) to produce 10-second samples of continuous vibrations excited by snakes, wasps, wind, rain, or routine embryo movements. Peak amplitudes from these continuous vibration samples were usually identical to peaks from longer samples and are not included separately. For embryo movements, we measured peak amplitudes from the 10-second continuous samples.

To test if attacks by different egg-predator species excited vibrations of different peak, RMS, and RMScont amplitudes, we performed Kruskal-Wallis tests. Because we did not record or sample particularly light wind and rain vibrations, we cannot statistically test whether these disturbances differ in amplitude from predator attack vibrations on average. We were, however, interested in whether some benign source disturbances excite vibration amplitudes similar to predator attack. We, therefore, used Mann Whitney *U* tests to compare the amplitudes of predator attack and benign source vibration samples.

### Amplitude-dependence of the hatching response: synthetic stimuli

To examine *A. callidryas*’ response to variation in the amplitude characteristics of vibrational stimuli, we conducted several playback experiments featuring vibration stimuli presented to either egg clutches or trays of individual embryos (Warkentin et al., 2006b; Warkentin et al., 2022).

To test whether *A. callidryas* respond to absolute variation in stimulus amplitude, and to determine if this is a graded or threshold response, we played synthetic vibrational noise to clutches of embryos using two different interfaces. For this first experiment, we played vibrational stimuli from a Macintosh G4 laptop computer through an Onkyo MSE-U33HB sound card and a custom-built amplifier (E. Hazen, Boston University Electronic Design Facility) to a Brüel and Kjær 4810 electryodynamic shaker. Attached to the shaker was the minishaker-clutch interface (MCI), a rigid stinger that terminates in a series of regularly spaced blunt metal tines. We mounted clutches on a rigid support stand, and carefully slid this stand forward, so that the tines of the MCI penetrated the clutch jelly, presenting both motion and contact cues. A shallow tray filled with aged tap water caught any hatched embryos. We used only clutches with ≥ 20 eggs. Five minutes following any hatching induced by set-up, we removed these hatched tadpoles and began playback. If ≥ 25% of a clutch hatched during set-up we did not use it for the experiment. We recorded the number of hatched embryos every minute during playback, and for five minutes thereafter. Following playback, we assessed the hatching competence of any remaining embryos through manual stimulation. Each clutch received a single stimulus presentation.

The base stimulus was constructed in Matlab as low-frequency filtered random white noise, played in a regular rhythmic temporal pattern previously shown to elicit high levels of escape hatching when played at an RMS amplitude of 4.4 m/s^2^ [0.5 second vibration duration, 1.0 second intervibration interval (Warkentin et al. 2006b)]. Intervals were created by setting values within specified periods to zero, leaving roughly rectangular amplitude envelopes of vibrational noise; all edits were made at zero-crossings to avoid introducing clicks. Each stimulus was five minutes in length. During June 2004, we played clutches stimuli at seven amplitudes (0.088, 0.132, 0.176, 0.22, 0.264, 0.352, and 0.44 m/s^2^ RMS, N = 79 clutches total). From these data it appeared as though induced hatching reached a maximum level beyond a certain amplitude. To test if hatching continued to increase at even higher amplitudes, we conducted an additional series of playback in June 2005. We presented clutches with seven stimuli (0.088, 0.22, 0.44, 0.66, 0.88, 1.32, and 1.76 m/s^2^ RMS, N = 95 clutches total). RMS amplitudes for the vibrational playback experiments conducted with simple synthetic stimuli, reported here and below, are most comparable to the RMScont amplitudes described above for recordings of natural disturbances (i.e. periods of silence between bursts of noise are excluded from RMS calculations).

For the 2005 playback series, we used an improved MCI (Warkentin et al. 2006b), designed to improve coupling with the clutch. In addition, stimuli for the 2005 experiment were equalized using custom scripts in MatLab (R13, MathWorks, Natick, USA) to correct for frequency filtering in our playback apparatus. These factors likely somewhat altered the playback stimulus, as perceived by embryos. Results from the playback series with the two MCIs are, therefore, considered separately.

To test for overall effects of amplitude variation on the hatching response and for effects of playback series, we used a general linear mixed model (GLMM) using binomial data (hatched vs. not hatched in clutch) and clutch as a random effect in R. We used Mann Whitney *U* tests to compare the hatching responses between the 2004 and 2005 series.

### Ontogenetic variation in the amplitude-dependence of hatching

To test how *A. callidryas* vibration sensitivity changes over development, we presented motion-only cues from synthetic vibrational noise to individual embryos using a tray interface and measured hatching responses across vibration amplitudes at three ages (4–6 days). The vibration playback system to present motion-only cues to trays of individual eggs was detailed elsewhere (Warkentin et al., 2019; Warkentin et al., 2022), but briefly, it consisted of an electrodynamic minishaker (LDS V203; Bruel and Kjær, Nærum, Denmark) interfaced with a custom-made amplifier (E. Hazen, Boston University Electronic Design Facility), which was connected via an external sound card (MSE-U33HB; Onkyo, Osaka, Japan) to a MacBook Air. Vibrational stimuli were played using the open-source program Audacity 2.1.0. We tested the frequency fidelity of vibration playbacks by recording playback stimuli using an accelerometer (AP32, AP Technology International, Oosterhout, The Netherlands) attached to a tray with plasticine. The accelerometer was connected via signal conditioner and external sound card (Onkyo MSE-U33HB) to a Macbook Pro laptop running Raven Pro 1.3 (Cornell Laboratory of Ornithology, Ithaca, NY). Based on frequency analysis of the recorded stimuli, we adjusted playback stimuli through Matlab to compensate for nonlinearities in the frequency response of the playback equipment (Warkentin et al., 2022). The base stimulus used in our playback experiments was constructed in Matlab as low-frequency filtered random white noise (0–100 Hz) and featured a regular rhythmic pattern of 0.5 s vibration duration and 1.5 s silence, which elicited the highest hatching in prior studies on temporal pattern effects (Jung et al., 2022; Warkentin et al., 2006b). Intervals were created by setting values within specified periods to zero, leaving roughly rectangular amplitude envelopes of vibrational noise. We also recorded the stimulus treatments with silent intervals removed to determine the RMS amplitudes in m/s^2^.

We performed three series of playback experiments, presenting the same vibration stimulus at different amplitudes to trays of up to 15 embryos across different stages of development. For the first series, conducted in 2016 between June 14 and June 29, we tested 163 trays of eggs in three different age groups: 4 d (at 19.69 ± 0.14 h, N = 55 trays), 5 d (at 16.78 ± 0.18 h, N = 60 trays), and 6 d (at 11.27 ± 0.17 h, N = 48 trays). Within each age group, all playbacks were conducted within 4 h of the mean time to reduce variation in hatching responses due to embryo development. Eggs were transferred to individual funnel-shaped spaces in egg trays at age 3 d, just before the embryos developed sensitivity to mechanosensory cues (Jung et al., 2019; Jung et al., 2020; Warkentin et al., 2017). For each trial, when eggs reached the desired age or developmental stage for testing, the trays were connected to the minishaker through a custom tray interface (Warkentin et al., 2022) to present vibrational cues. After attaching the tray to the shaker’s interface, we allowed 5 minutes of undisturbed acclimation before starting a playback. We noted any hatching induced by the setup procedure and only used trays with at least 5 eggs remaining after 5 minutes of acclimation. We presented the vibration stimulus to embryos in each of the three age groups at four amplitudes in randomized order: 2 m/s^2^ (N = 40 trays), 4 m/s^2^ (N = 41 trays), 8 m/s^2^ (N = 40 trays), and 15 (N = 42 trays) m/s^2^. We simultaneously started the vibration playback and the timer to record hatching events, then watched the eggs closely through the playback period to record the latency for the first embryo in the tray to hatch.

Hatchlings slid through the funnel and into a container of aged tap water below the tray, allowing us to keep a precise record of hatching latencies for individuals. After the end of the playback, we watched the eggs for 3 additional minutes and noted any additional hatching, then manually stimulated any unhatched embryos to test for hatching competence. The number of embryos that hatched in response to playback and the number of healthy and hatching-competent embryos that did not hatch were counted at the end of each experiment. These counts were used to calculate the final proportion hatched. Any embryos that failed to hatch in response to manual tactile stimulation were excluded from calculations (Jung et al., 2022; Warkentin et al., 2019). Most clutches contributed eggs to fill multiple trays, but each tray was used for only one trial.

The following two playback series followed the same experimental procedure as the first, but with only one intermediate age group using successively lower stimulus amplitudes. The second series was conducted between June 21 and August 6, 2016. We followed the same experimental procedure using 119 trays of eggs at age 5 d (at 17.08 ± 0.13 h). Amplitudes were 0.15 m/s^2^ (N = 30 trays), 0.3 m/s^2^ (N = 30 trays), 0.8 m/s^2^ (N = 30 trays), and 2 m/s^2^ (N = 29 trays). Our final series included playbacks to 104 trays of eggs at 5 d (at 18.68 ± 0.32 h) with amplitudes of 0.05 m/s^2^ (N = 29 trays), 0.09 m/s^2^ (N = 17 trays), 0.12 m/s^2^ (N = 14 trays), 0.15 m/s^2^ (N = 16 trays), 0.6 m/s^2^ (N = 14 trays), and 2 m/s^2^ (N = 14 trays). The third series with the lowest amplitude stimuli was conducted between July 7 and July 20, 2017. Each stimulus set included overlapping amplitudes to facilitate comparisons across series and test for potential effects of time in the season. Since stimulus sets were created and played the same way in this series, using the same equipment across 2016 and 2017, and hatching responses were consistent across the series for the amplitude played across all three sets (GLMM, *χ*^2^ = 1.2421, *df* = 2, *P* = 0.5274), we pooled data across the three sets. To evaluate the effects of stimulus amplitude and embryo development on hatching response (binomial data) and hatching latencies (Poisson data), we employed generalized linear mixed models with clutch as a random effect.

### Amplitude-dependence of the hatching response: natural stimuli

To assess if the effects of amplitude interact with other stimulus properties, and, specifically, to determine how amplitude variation in natural egg-clutch disturbances affects escape hatching, we played clutches recorded rain and snake attacks at varying amplitudes. Stimuli included three recorded exemplars of rainstorm vibrations (205 s, 205 s, and 240 s in length) and two snake attack (*L. ahaetulla*) stimuli. One snake stimulus consisted of a complete uninterrupted attack (snake 1, 240 s, referred to as ‘single’ in Warkentin 2005). The other snake stimulus was a composite of three separate *L. ahaetulla* attacks, with periods of accelerometer biting edited out, spliced together to form one 205 s stimulus (snake comp, referred to as ‘composite’ in Warkentin 2005). In previous fixed amplitude playbacks, these snake exemplars elicited substantially more hatching than these rain exemplars, consistent with relative levels of hatching seen in response to each disturbance type in nature (Warkentin 2005). For the experiment presented here, we adjusted the RMS amplitude of each exemplar to approximately 0.17 ms^-2^ (0.17 ± 0.049 SD; range, 0.11–0.25 m/s^2^) to produce a base (1X) stimulus. Natural disturbance vibrations are very dynamic and include considerable amplitude modulation. Unlike for our synthetic stimuli, RMS amplitude calculations for natural stimuli included any brief periods of relative silence. It is not known how *A. callidryas* perceives the amplitude properties of a vibrational disturbance. It is, therefore, not clear whether Peak, RMS, or some other measure of amplitude is most appropriate for studying the defensive behavior of these embryos. Rain vibrations have higher peaks relative to their RMS amplitude than do snake attacks. Thus, it was not possible to match rain and snake attack exemplars in both RMS and peak amplitude. Because we were interested in comparing embryos response to amplitude variation of snake attack and rainstorm vibration, rather than responses at any particular amplitude, we chose base stimulus amplitudes for each exemplar that would provide a large range of overlap in both RMS and peak amplitude over the range across which they were amplified. We presented all clutches with either this base stimulus, or a stimulus three or five times the amplitude of the base stimulus. Thus, all exemplars were played at 1X, 3X, and 5X base levels. For one rain and one snake exemplar, we played clutches five additional amplitudes as follows: 0.5X, 2X, 4X, 6X, and 7X base stimulus amplitude. All stimuli were equalized for playback using MatLab, and the frequency fidelity of playback was checked via rerecording. Data were positively skewed and fit a negative binomial distribution and log link function. Therefore, we tested the effects of disturbance type, exemplar, playback amplitude, and disturbance type by amplitude interaction on the clutch-size-adjusted number of embryos hatched from each clutch using a generalized linear model (GzLM) in SPSS (v.16, SPSS Inc., Chicago, USA).

### Influence of amplitude modulation on the hatching response

To test whether embryos use the fine-scale amplitude modulation of disturbance vibrations to inform their hatching decision, we played synthetic stimuli matched in global amplitude and duty cycle, but varying in amplitude envelope shape. Stimuli were bursts of 0–100 Hz band limited white noise in a one second noise, one second inter-vibration interval temporal pattern, played at 0.48 m/s^2^ RMS. Each stimulus was five minutes in length. We played three stimuli. In the first stimulus (‘fade in’), each burst of noise steadily increased in amplitude throughout its 1 s duration and ended abruptly (N = 20 clutches). In the second stimulus (‘diamond’), bursts of noise increased more rapidly, reached peak amplitude at mid-duration, and then decreased in amplitude, reaching zero amplitude at 1 s in duration (N = 20). The third stimulus (‘fade out’) was identical to the fade in stimulus, but was played in reverse, such that each noise burst started at full amplitude and decreased throughout its 1 s duration (N = 20). To test whether amplitude envelope shape affected the proportion of embryos in a clutch that hatched, we used a Kruskal-Wallis test, and post hoc Dunn test for pairwise comparisons.

## Results

### Amplitude characteristics of common natural vibrational disturbances to egg clutches

Each type of vibrational disturbance to *A. callidryas* egg clutches varied widely between recordings in all three measures of amplitude (Fig. 1, Kruskal-Wallis: peak, *χ*^2^ = 55.47, *df* = 7, *P* = 1.2e-9; RMS, *χ*^2^ = 33.82, *df* = 6, *P* = 7.3e-6, RMScont, *χ*^2^ = 39.901, *df* = 7, *P* = 1.3e-6). There was also a considerable amount of variation between attacks from different predator species and between different benign disturbances in at least two measures of amplitude (Kruskal-Wallis: RMS_predator_, *χ*^2^ = 22.11, *df* = 4, *P* = 0.00019; RMScont_predator_, *χ*^2^ = 14.73, *df* = 4, *P* = 0.0053, peak_non-predator_, *χ*^2^ = 28.76, *df*=2, *P* = 5.693e-7; RMScont_non-predator_, *χ*^2^ = 22.55, *df* = 2, *P* = 1.267e-5). For predators, wasp and *L. ornata* attacks elicited vibrations of lower RMS amplitude than did attacks by the other predators in all tested measures of amplitude (Kruskal-Wallis: peak, *χ*^2^ = 6.07, *df* = 1, *P* = 0.01374; RMS, *χ*^2^ = 21.91, *df* = 1, *P* = 2.851e-6, RMScont, *χ*^2^ = 12.69, *df* = 1, *P* = 0.0003681). For benign disturbances, embryo movements excited much lower amplitude vibrations than wind and rain in all tested measures of amplitude (Kruskal-Wallis: peak, *χ*^2^ = 19.85, *df* = 1, *P* = 8.379e-6; RMScont, *χ*^2^ = 22.35, *df* = 1, *P* = 2.278e-6).

**Figure 1.**
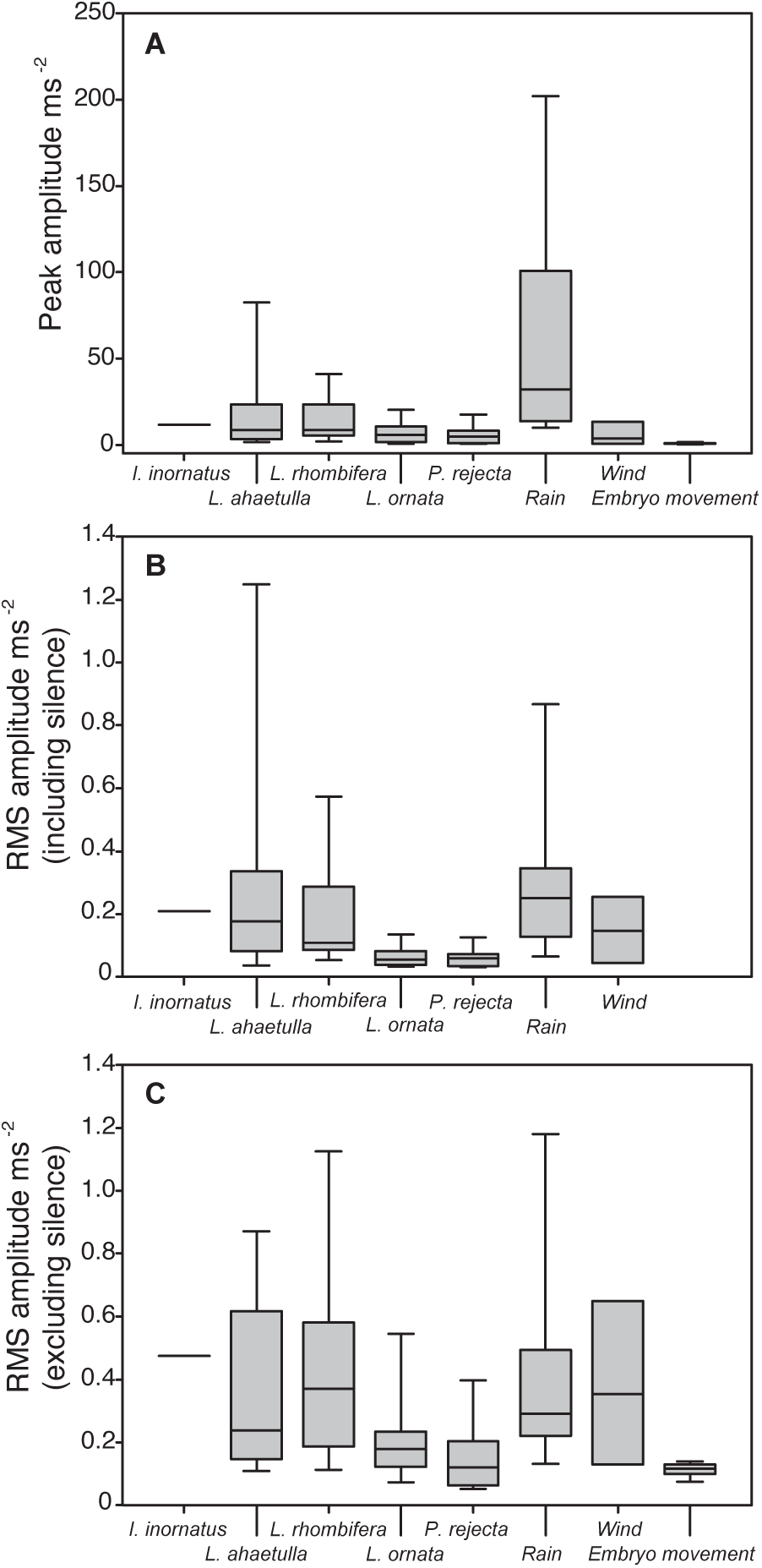
Amplitude characteristics of common natural vibrational disturbances to *Agalychnis callidryas* egg clutches reveal that no disturbance amplitude was unambiguously indicative of predator attack. Shown are median, 10^th^, 25^th^, 75^th^, and 90^th^ percentiles for snake attacks (*Imantodes inornatus, Leptophis ahaetulla, Leptodeira rhombifera,* and *Leptodeira ornata*), wasp attacks (*Polybia rejecta*), and vibrations excited by hard rain, strong wind, and routine embryo movements. (**A)** Peak amplitudes for long periods of each disturbance type. (**B)** RMS amplitudes for long periods of each disturbance type, including intermittent periods of silence. RMS amplitude for embryo movements was not measured separately for long period samples. (**C)** RMS amplitudes for 10 s periods of continuous vibrations sampled from each disturbance, constructed from longer samples by editing out periods of silence. Hard rain is the highest amplitude disturbance. Percentiles are not shown for *I. inornatus* attacks, and 10^th^ and 90^th^ percentiles are not shown for wind recordings, due to low sample sizes.

Perhaps of most relevance to *A. callidyas’* escape hatching decision, when all common disturbance types were considered together, there was a large amount of overlap between the amplitudes excited by predator attacks and those excited by benign sources (Fig. 1). Predator attacks, considered as a group, did not differ from benign source vibrations in overall peak amplitude or the RMS amplitude of continuous vibration samples (Mann Whitney: peak, *Z* = 1.20, *P* = 0.8841; RMScont, *Z* = 1.70, *P* = 0.96). The long-period RMS amplitudes of predator attacks were statistically lower than those of long periods of benign disturbance (RMS, *Z* = 3.01, *P* = 0.0013). This difference, however, was driven by the fact that long-period RMS amplitudes for embryo movements were not measured and long-period vibration samples from hard rain were significantly higher in all amplitude measures than predator attacks (Mann Whitney: peak, *Z* = 4.96, *P* = 3.613e-7; RMS, *Z* = 3.32, *P* = 0.0004464, RMScont, *Z* = 2.17, *P* = 0.01511).

### Amplitude-dependence of the hatching response: synthetic stimuli

The number of 5 d old embryos hatching in response to playback of synthetic noise stimuli varied with stimulus amplitude in both series conducted (Fig. 2, GLMM, *χ*^2^ = 210.05, *df* = 10, *P* < 2e-16), but we did not find an interaction between the effects of amplitude and series (GLMM, *χ*^2^ = 0.10, *df* = 2, *P* = 0.9519). Embryo response differed between the series conducted with the original (2004) and the improved (2005) MCI (Mann Whitney: peak, *Z* = 2.65, *P* = 0.00397), but still the number of embryos hatching in response to playback of synthetic noise increased with stimulus amplitude in each separate series (GLMM_2004_, *χ*^2^ = 74.38, *df* = 6, *P* = 5.139e-14; GLMM_2005_, *χ*^2^ = 131.69, *df* = 6, *P* < 2.2e-16). Within each series, hatching increases in a nearly linear relationship with stimulus amplitude for a portion of the response curve and then appears to reach a threshold, above which further increases in stimulus amplitude are not associated with corresponding increases in hatching response (Fig. 2). This apparent plateau of maximal hatching for each series is not explained by a simple logarithmic or power relationship between stimulus amplitude and embryo response; a distinct plateau remains if results are plotted on a log scale. All clutches tested in these assays were 5 days post oviposition.

**Figure 2.**
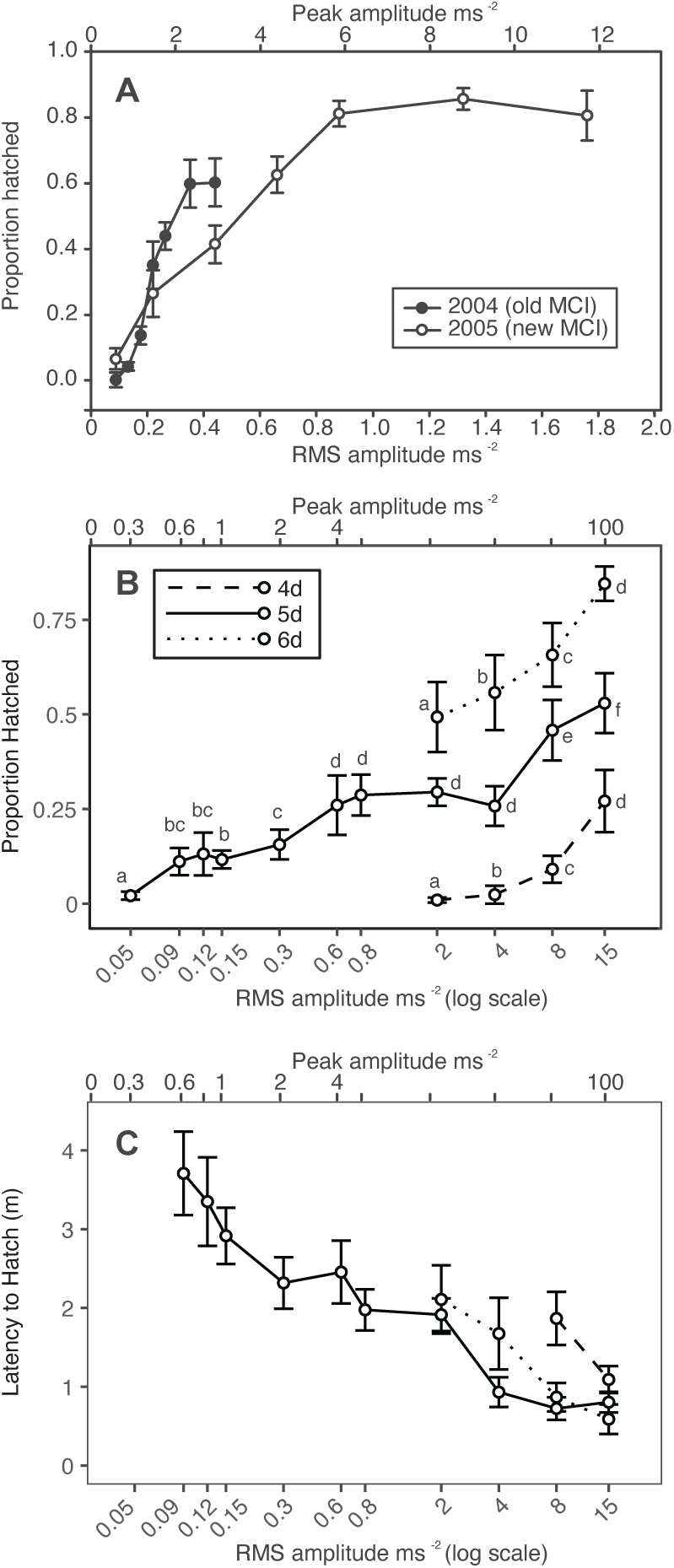
The hatching response of *Agalychnis callidryas* embryos to playbacks of vibrational noise at varying amplitudes. **(A)** Two playback series were conducted using bands of low-frequency noise in a 0.5 s noise/1 s silence temporal pattern. The tine playback apparatus – which presents both motion and tine-contact cues to entire egg clutches – was altered between the two series, both presented to 5-day-old embryos. Escape hatching increases with amplitude until a plateau of maximal hatching is reached. Shown are mean proportion of embryos within each clutch that hatched, ± SE, for each stimulus presented. **(B, C)** Ontogenetic variation in the amplitude-dependence of hatching using bands of low-frequency noise in a 0.5 s noise/1.5 s silence temporal pattern. All stimuli were played through a tray apparatus designed to present vibrations without contact cues to a set of embryos held individually. Escape hatching increases with both amplitude and development. Means not sharing any letter within age groups are significantly different by a post-hoc test at the 5% level of significance. Shown are **(B)** mean proportion of 4, 5, and 6 d old embryos in trays that hatched, ± SE, for each stimulus presented, and **(C)** the latency of the first embryo in each tray to hatch in response to vibration stimuli of varying amplitudes.

### Ontogenetic variation in the amplitude-dependence of hatching

Vibration playback experiments using trays of individual embryos across a broader range of ontogeny (4-, 5-, and 6-days post oviposition) revealed that hatching increased with embryo development (GLMM, *χ*^2^ = 821.97, *df* = 2, *P* < 2.2e-16) and stimulus amplitude (χ^2^ = 1892.19, *df* = 10, *P* < 2.2e-16), with an interaction effect between the two (χ^2^ = 124.48, *df* = 6, *P* < 2.2e-16, Fig. 2B). The number of embryos hatching in response to playback of synthetic noise varied with stimulus amplitude in each individual age group (GLMM_4 days_, *χ*^2^ = 450.35, *df* = 3, *P* < 2.2e-16; GLMM_5 days_, *χ*^2^ = 1458.9, *df* = 10, *P* < 2.2e-16; GLMM_6 days_, *χ*^2^ = 60.579, *df* = 3, *P* < 4.421e-13).

For each age group, hatching increased in a nearly linear relationship with stimulus amplitude for a portion of the response curve, with no evidence of an upper threshold where the response stabilizes; hatching increased in response to 15 ms^2^ relative to 8 ms^2^ in all age groups tested (Fig. 2B). This contrasts with playback experiments to clutches, where tines present both motion and contact cues, wherein above a high-amplitude threshold further increases in stimulus amplitude are not associated with corresponding increases in hatching response.

At 5 days, 2 ± 5% of embryos hatched at the lowest playback amplitude we tested, 0.05 m/s^2^, and their response was significantly higher at 0.09 m/s^2^ (11 ± 14%, Fig. 2B). In contrast, less than 1% of younger, 4-day-old embryos hatched at 2 m/s^2^. Their response increased significantly, to 2 ± 9% at 4 m/s^2^, and then to 9 ± 12% at 8 m/s^2^ (Fig. 2B). Thus, whether we consider 2% or near 10% as a “threshold” hatching response, older embryos responded to much lower amplitude vibrations, at least 80-fold lower.

Stimulus amplitude affected the latency to hatch across all ages tested (GLMM, *χ*^2^ = 1328.08, *df* = 10, *P* < 2.2e-16, Fig. 2C). Embryo age also influenced hatching latency (GLMM, *χ*^2^ = 80.02, *df* = 2, *P* < 2.2e-16), and there was an interaction effect between age and amplitude (GLMM, *χ*^2^ = 63.32, *df* = 6, *P* < 9.515e-12). Latency to hatch varied with stimulus amplitude in each individual age group (GLMM_4 days_, *χ*^2^ =18.83, *df* = 1, *P* < 1.431e-5; GLMM_5 days_, *χ*^2^ = 1012.70, *df* = 9, *P* < 2.2e-16; GLMM_6 days_, *χ*^2^ = 335.53, *df* = 3, *P* < 2.2e-16).

### Amplitude-dependence of the hatching response: natural stimuli

Hatching increased with amplitude for recorded rain and snake attack exemplars as well (Fig. 3, GzLM, *χ*^2^ = 60.60, *df* = 7, *P* < 0.001). The amplitude-dependent response to rain and snake attack vibrations differed, however. Hatching increased more steeply as a function of snake vibration amplitude than it did as a function of rain vibration amplitude, as reflected in the significant interaction between the effects of vibration amplitude and disturbance type (GzLM, *χ*^2^ = 51.0, *df* = 7, *P* < 0.001). After accounting for the effects of disturbance type, there was also a borderline significant effect of individual rain or snake attack exemplar (GzLM, *χ*^2^ = 7.57, *df* = 3, *P* = 0.056). In at least one of the two snake attack exemplars (snake1), hatching did not increase beyond a certain amplitude, resembling the responses we observed to playback of synthetic stimuli.

**Figure 3.**
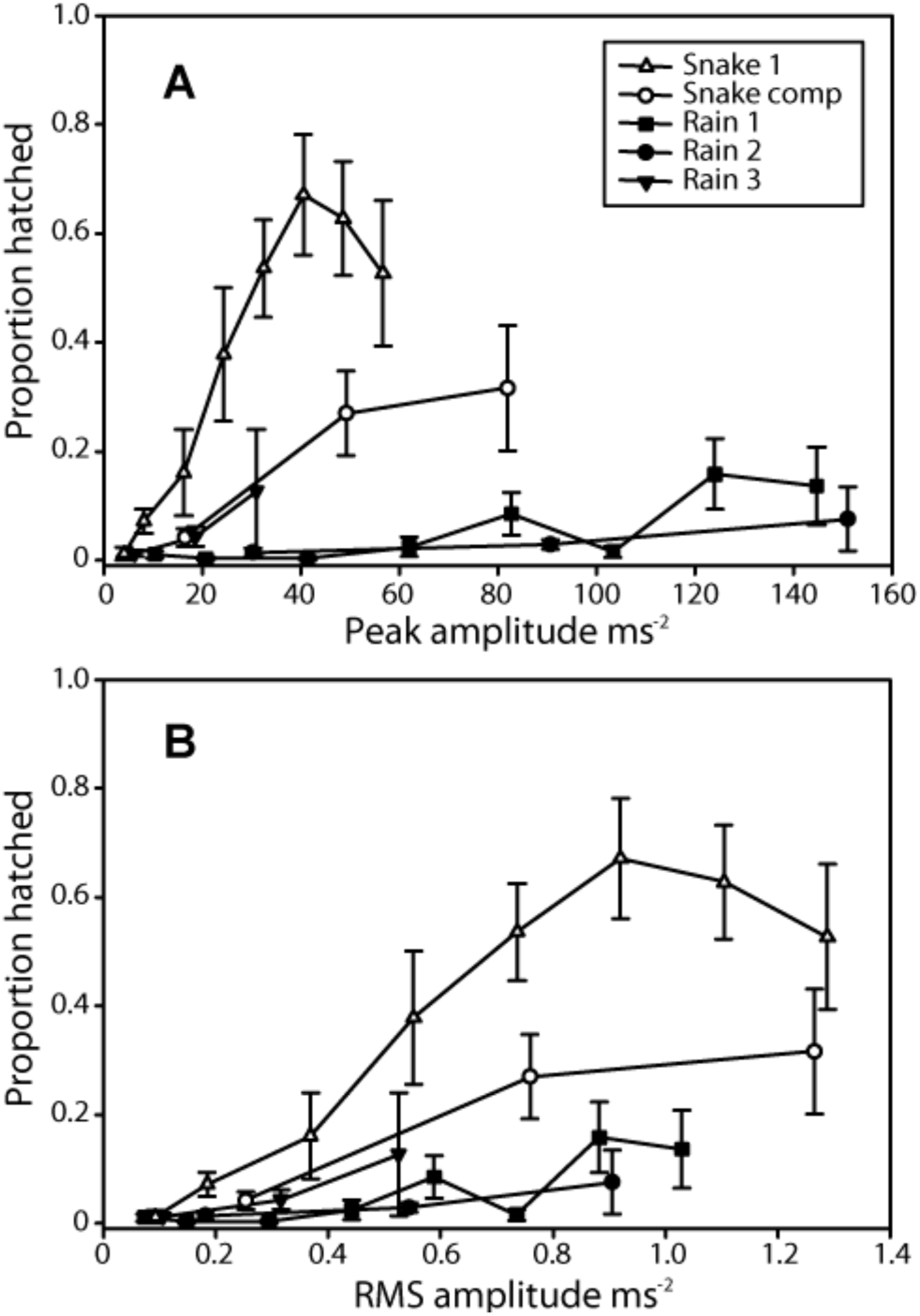
The hatching response of *Agalychnis callidryas* embryos to playbacks of recorded snake attack (*Leptophis ahaetulla*) and rainstorm exemplars at varying amplitudes. Embryos were presented with two snake attack and three rain exemplars. One snake and one rain exemplar were presented at eight amplitudes, the remaining exemplars were presented at three amplitudes. Hatching increases more strongly as function of amplitude for snake attacks than it does for rainstorm vibrations. **A)** Hatching response as a function of peak amplitude. **B)** Hatching response as a function of RMS amplitude. Data are mean proportion hatched ± SE for each stimulus presented.

### Influence of amplitude modulation on the hatching response

The hatching response of *A. callidryas* embryos also varied with the amplitude envelope shape of synthetic stimuli (Fig. 4, Kruskal-Wallis, *χ*^2^ = 7.49, *df* = 2, *P* = 0.024). The ‘fade in’ stimulus elicited less hatching than either the ‘fade out’ (Dunn, *Z* = 2.26, *P* = 0.0239) or the ‘diamond’ (*Z* = 108.5, *P* = 0.0136) stimuli. We found no difference between the response of embryos to the ‘diamond’ and ‘fade out’ stimuli (*Z* = 0.21, *P* = 0.835).

**Figure 4.**
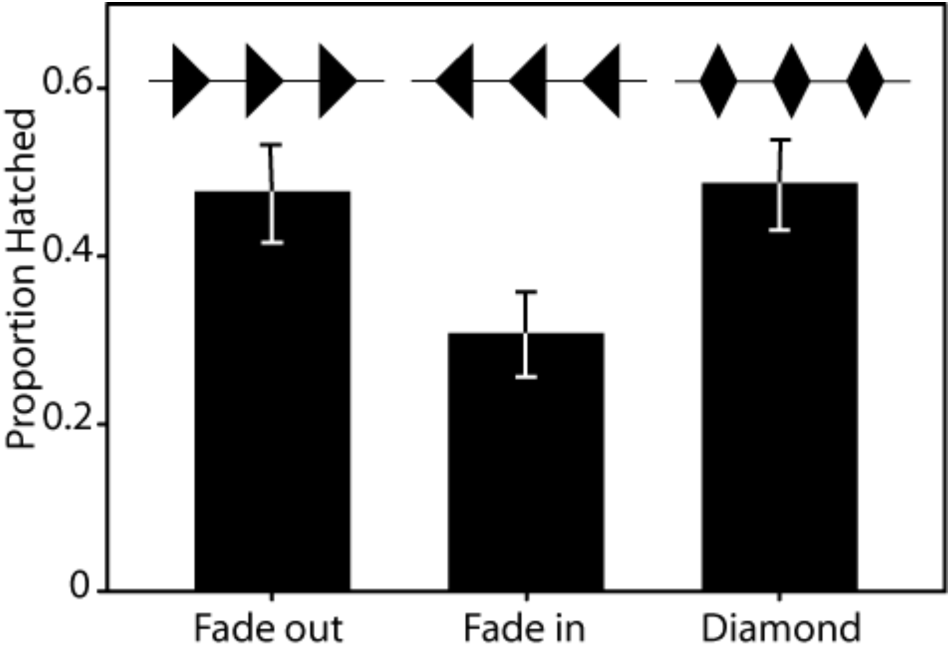
The hatching response of *Agalychnis callidryas* embryos to playbacks of synthetic vibrational stimuli varying in amplitude envelope shape. Stimuli were matched for frequency (0–100 Hz band limited white noise), temporal pattern (1 s noise/1 s silence), and peak and RMS amplitude, but varied in rise and fall times as shown in the waveform graphics above the data. ‘Fade in’ elicited less hatching than the other two stimuli. Shown are mean proportion hatched ± SE for each stimulus.

## Discussion

*Agalychnis callidryas* embryos respond in a graded manner to amplitude variation in vibrational disturbances to their egg-clutches. For playbacks that combine motion with contact cues, the range over which hatching varies with amplitude is consistent with the amplitude range of predator attacks, and embryos appear to reach a plateau of maximal hatching at RMS amplitudes above the range commonly experienced during natural disturbances (Fig. 1, 2A). Clearly, embryos use amplitude information to refine their hatching decision. Embryos are more likely to hatch and hatch faster as they get older (Fig. 2B, C). However, the absolute amplitude characteristics of vibrational disturbances to *A. callidryas* egg-clutches do not reliably predict predation risk.

Predator disturbances and strong benign-source disturbances share similar broad amplitude ranges (Fig. 1). However, at matched amplitudes, embryos show a stronger response when disturbed by vibrations from snakes than those from rain. Thus, the nature of *A. callidryas’* response to disturbance amplitude depends other vibrational properties assessed for the escape hatching decision, and this interaction is consistent with relative threats posed by snake attack and rain disturbances. Embryos also respond to fine-scale modulation in the amplitude of clutch vibrations (Fig. 4); however, further work is necessary to determine whether amplitude envelope characteristics improve *A. callidryas’* ability to discriminate between dangerous and benign vibrational disturbances.

### Amplitude characteristics of common clutch disturbances

Embryos experience a range of common clutch disturbances that vary greatly in amplitude characteristics. To examine just how loud these disturbances are in nature, we recorded several natural disturbance types. We found there was a great deal of amplitude variation in each type of common clutch disturbance, with the exception of routine embryo movements, and that the range of amplitudes excited by predator attacks broadly overlaps that excited by benign sources (Fig. 1). There is no peak or RMS amplitude of either full disturbances or edited periods of continuous vibration that is unambiguously characteristic of predator attack. Hard rain, in fact, excited the highest amplitude vibrations for all measures. Strong wind excited high amplitude vibrations as well. While one might presume that the highest amplitude stimuli should yield the highest hatching, our recordings indicate that this is not the case, and the highest intensity stimulus is actually a non-threatening one. Although we deliberately excluded very low amplitude wind and rain recordings from our analyses, the exemplars chosen were within the range commonly experienced by clutches at our field site in Gamboa.

The wasp *Polybia rejecta* and the snake *Leptodeira ornata* both excited vibrations that were of lower RMS amplitude than the three other egg predators. In the case of *P. rejecta*, this could be due to the wasp’s small size. The idea that disturbances by smaller, less massive sources generate lower amplitude vibrations is consistent with published patterns of reduced amplitude for a larger vs. small water drop (Güell et al., 2024). The relatively low amplitude vibrations produced by *L. ornata* snakes are more puzzling, especially considering that the congeneric *L. rhombifera* excited much higher amplitude vibrations. The feeding behavior of *L. ornata* is, however, distinct from that of the other snake species examined here (D’Amato and Warkentin, unpublished). *L. ornata* feeds more efficiently, making fewer movements in contact with eggs that do not result in egg ingestion. Also, its attacks consist of a few, relatively long periods of contact with the clutch, rather than many short bites. If strikes and tearing off mouthfuls of eggs generate more intense vibrations than do pterygoid walks through a clutch, this could contribute to the difference in disturbance amplitudes excited by different snake species.

Interestingly, the range of amplitude-dependence of hatching in response to a synthetic stimulus (Fig. 2A) and two *L. ahaetulla* exemplars (Fig. 3) is relatively high compared with the amplitudes of recorded *L. ornata* and *P. rejecta* attacks (Fig. 1). Although this comparison may suggest that escape hatching might be less effective in response to the more vibrationally subtle predators, that is not the case. Escape success of fully hatching-competent embryos is ∼80% in attacks by both *P. rejecta* and *L. ornata*, similar to that found in the vibrationally higher amplitude attacks by *L. rhombifera* and *L. ahaetulla* (Gomez-Mestre and Warkentin, 2007; Warkentin et al., 2006a) (also D’Amato and Warkentin unpublished). Although hatching responses varied with each stimulus presented, overall, at equivalent amplitudes, embryo response was lower to playback of recorded snake attack exemplars than to attack by live snakes. This may reflect limitations in the fidelity of our playback apparatus or may be the result of background noise present in our recordings. Alternatively, it may indicate that embryos also respond to non-vibrational predation cues that are not reproduced by our playbacks, such as light (Güell and Warkentin, 2018) or elements of tactile stimulation (Fouilloux et al., 2019).

### Graded vs. categorical defensive response to amplitude variation

The magnitude of *A. callidryas’* hatching response varied gradually over a wide range of stimulus amplitudes or embryo ages, for both synthetic stimuli and for rain and snake exemplars (Figs 2, 3). Furthermore, incremental increases in hatching were nearly linear over much of this range, including over 10 dB ranges of the 2005 synthetic noise series (Fig. 2A), the developmental series (Fig. 2B, C), and the ‘snake1’ exemplar (Fig. 3). The presence of this graded response is strong evidence that variation in the amplitude of vibrational disturbances is informative to embryos making their hatching decision.

Interestingly, some of the behavioral response curves (i.e. both synthetic series to clutches, snake1) seem to reach a plateau of maximal hatching at some threshold amplitude (Fig. 2A, Fig. 3). While clutches are bound by an upper limit of hatching (100%), the plateau occurred at a different response level for each stimulus, thus was not likely due to a boundary effect. Thus, at least for tine playbacks presenting motion and contact cues, there seems to be an upper amplitude threshold, which depends on various stimulus properties, above which embryos do not hatch more with additional increases within the range of natural amplitude variation.

During some tray-based (motion-only) playbacks, we played stimuli at amplitudes well in excess of those observed for common clutch disturbances in nature (Fig. 2B). At these high amplitudes, hatching continued to increase with stimulus intensity, without ever leveling off. Likewise, a previous experiment, utilizing similarly high amplitude stimuli, consistently elicited nearly complete hatching from egg clutches of the same developmental maturity, demonstrating that 100% hatching is possible in tine playbacks (Caldwell et al., 2009). It remains to be seen, however, whether amplitude-dependent variation in hatching at these high amplitudes has any adaptive significance.

Using a behavioral criterion of either 2% or near 10% hatching to indicate responsiveness, the lower amplitude threshold for hatching decreased substantially (ca. 80-fold) with development. While just 9% of 4-day-old embryos hatched in response to a synthetic stimulus played at 8 m/s^2^ RMS, the same stimulus played at 0.09 m/s^2^ elicited 11% hatching at age 5 days (Fig. 2B). Thus, the older embryos responded to much lower-amplitude vibrations. We know that later in embryo development, decision rules change in ways that reflects the developmental decrease in larval-stage risk (Caldwell et al., 2010; Jung et al., 2021; Jung et al., 2022; Warkentin et al., 2019). However, at the onset of mechanosensory-cued hatching, increasing sensory ability is likely key (Jung et al., 2019; Jung et al., 2020). In nature, relatively few embryos at 4 days of age hatch during wasp attacks; most are taken by wasps and presumably killed. By contrast most 5-day-old embryos escape (Gomez et al., 2023). Considering the low amplitude of vibrations induced by wasp attacks (Fig. 1), this change in fates fits well with the amplitude response curves we present here (Fig. 2B). Moreover, the relatively high threshold for 4- day-old embryos to respond to a motion-only cue (Fig. 2B) supports that more complex mechanosensory cues (i.e. direct contact cues) are important for the embryo escape from wasps that does occur at that age. 5-day-old embryos hatch when wasps attack other eggs, transmitting vibration cues with no direct contact, whereas 4-day-old embryos respond only to direct attacks on their own egg, which provide more complex cues (Gomez et al., 2023). It is important to note that while we saw very little hatching in response to our lowest amplitude stimuli, we cannot conclude that our behavioral response curves represent the neurophysiological limits of *A. callidryas*’ sensitivity to vibration. Embryos may detect lower amplitude vibrations, but not respond to them (Ydenberg and Dill, 1986).

### Does amplitude information convey the immediacy of predation threat?

Peak and RMS amplitudes of vibrations excited by benign sources and predator attack did not consistently differ, and amplitude information may not, therefore, be useful when discriminating between disturbance types. Still, amplitude variation could provide information about the immediacy of a predator threat. This would clearly be valuable during wasp attacks, as wasps can only take one egg at a time and hatching tends to be localized within clutches, near where the wasp is biting (Hughey et al., 2015). It also seems relevant in some snake attacks, particularly for species that feed more slowly, consuming eggs one by one (e.g. *L. rhombifera, Imantodes inornatus)*. It is not uncommon for both snakes and wasps to leave partially consumed clutches (Warkentin, 1995; Warkentin, 2000; Warkentin et al., 2006a), thus attack vibrations do not necessarily mean imminent death for embryos. An embryo may experience relatively low amplitude vibrations while a predator is extracting a sibling from the other side of the clutch. Higher amplitude vibrations may indicate a closer, and therefore more immediate, threat.

There has been little work on the estimation of distance to a signal source (‘ranging’) using amplitude characteristics of incidental predator cues. Some receivers do use attenuation and amplitude-dependent degradation of communication signals as ranging cues, however. This is commonly observed as ‘passive choice’ in the context of mate assessment, where the receiver shows a preference for what is perceived as the higher amplitude, and thus closer of multiple signals (Caldwell et al., 2022; Parker, 1982; Searcy and Andersson, 1986). Although passive choice involves a relative assessment of distance, there is evidence that some receivers can also assess the absolute distance to a signal source based on passive amplitude cues (Mershon and King, 1975; Nelson, 2000). Such estimates of absolute distance would likely be more useful in the assessment of predation risk.

In biomechanical testing of *A. callidryas* clutches, vibration amplitude decreases with distance from a pendulum impact, suggesting these embryos could use attenuation to assess distance (Güell et al., 2024). There is some question as to how useful amplitude properties of substrate vibrations are as a potential distance cue, especially for those that propagate through complex organic media. Unlike the majority of pressure waves in air and water (Bradbury and Vehrencamp, 1998), plant-borne vibrations do not decrease monotonically in amplitude as a function of distance from a vibration source (Michelsen et al., 1982). While the vibrational mechanics of *A. callidryas* egg clutches appear to behave non-linearly with respect to attenuation of energy at some frequencies, some frequencies present in wasp vibrations do attenuate predictably as they propagate through a clutch (Caldwell, 2010).

### Interactions between vibration amplitude and information from other properties

Despite different researchers conducting these experiments – decades apart – using different methods and different equipment, we recorded some similar patterns of hatching response to a range of amplitudes. The range of RMS amplitudes that induced hatching in all three playback setups (trays and two versions of the MCI) were comparable (Fig. 2). While hatching increased with amplitude for each stimulus presented during this study, the slope of the increase varied with stimulus. Differences between the MCI and frequency equalization used for the 2004 and 2005 synthetic noise series likely account for the difference in response we see between them. Interestingly, at the same age (5 days) and comparable amplitudes, overall hatching responses were greater during full-clutch MCI playbacks than they were during tray playbacks to separated embryos (Fig. 2), even though prior work (Warkentin et al. 2006b) found the temporal pattern used in the tine playbacks elicited less hatching than the pattern used in the tray playbacks (0.5 s noise, 1 s silence, 60% hatched vs. 0.5 s noise, 1.5 s silence, 74% hatched). We suspect that the greater response to tine playbacks is largely due to tactile and pressure cues introduced by the tines of the MCI as they move within the clutch jelly, bumping into and temporarily deforming eggs, and the consequent stimulation of the lateral line as well as vestibular system mechanoreceptors (Jung et al., 2019; Jung et al., 2020).

Of particular importance is the dramatic difference in amplitude-dependent response we see to snake and rain stimuli. The hatching response to playback of snake exemplars increased rapidly with increases in presentation amplitude. Despite the fact that rain vibrations excite the entire range of frequencies found in predator attacks (Caldwell et al., 2009), hatching responses to playbacks of rain exemplars were substantially lower and the increase in hatching in response to increases in amplitude were much smaller. Over the range of RMS amplitudes tested, hatching increases at a rate approximately 3-fold higher for snake attack than it does for rain vibrations. The rate of increase is approximately 6.5-fold higher if peak amplitudes are considered. Stimulus-specific variation in the amplitude-response of escape hatching may explain the strong response to some relatively low amplitude wasp and *L. ornata* attacks in nature (Gomez-Mestre and Warkentin, 2007; Warkentin et al., 2006a).

Differences between the amplitude-dependent response to playback of rain disturbance and snake attack vibrations must be due to non-amplitude properties of each disturbance type. We know that, for their hatching decision, *A. callidryas* non-redundantly combine information about the duration of individual vibrational events, the spacing between these events, the presence of low frequency energy characteristic of predator attack, as well as the absence of high frequency energy and initial periods of low intensity vibrations characteristic of rainstorms (Caldwell et al., 2009; Caldwell et al., 2010; Warkentin et al., 2006b). The results presented here indicate that vibration amplitude information influences the hatching decision as well, and that it is integrated along with these other predictors of risk.

### Response to amplitude modulation

Embryos may also integrate information about the fine-scale amplitude modulation of vibrational disturbances into this complex hatching decision. Our results clearly demonstrate that embryos respond to variation in the amplitude envelope of short bursts of noise, showing less hatching in response to the ‘fade in’ stimulus (1 s rise time) than to the two other stimuli presented. It is not clear, however, if this specific response is adaptive or relevant to defensive decisions made in nature. Vertebrates commonly show less dramatic defensive behavior or higher response thresholds to stimuli that rise more slowly in amplitude (Davis, 1984). Whether this is an evolved response or a byproduct of prey neurophysiology is not well understood. Still, in the case of *A. callidryas*, reduced hatching to stimuli with relatively long rise times could be adaptive. Many snake attacks on *A. callidryas* clutches are characterized by high amplitude strikes following periods of relative vibrational silence. Rainstorms, by contrast, have a tendency to build in amplitude over time (Caldwell et al., 2010). Past work has shown that embryos use some aspect (amplitude, frequency, etc.) of larger scale (60–180 s) periods of increasing intensity vibrations found at the start of rainstorms to avoid hatching to these benign stimuli (Caldwell et al., 2010). Embryos might use features of finer-scale amplitude modulation to distinguish snake attacks as well. In light of this possibility, it is interesting that we observed no difference in hatching responses to the abruptly starting ‘fade out’ stimulus and the ‘diamond’ stimulus, with an intermediate rise time (0.5 s). Further playback experiments as well as a more thorough description of the amplitude modulation characteristic of each common disturbance to *A. callidryas* egg-clutches will be necessary to understand how and why embryos incorporate information about fine-scale amplitude modulation into their hatching decision.

### Conclusion

The adaptive significance of any behavior is dependent upon the environment in which it naturally occurs. By studying the escape-hatching response of *A. callidryas* in the context of the vibrational disturbances that are relevant to this behavior in nature we have a better opportunity to uncover the properties of those disturbances that are informative to embryos, and to understand how these are used to make defensive decisions in face of possible costly decision errors. *A. callidryas* embryos modulate escape hatching in response to variation in the amplitude properties of vibrational disturbances. Embryos, however, do not base their decisions on disturbance amplitude alone, nor does it appear that such a strategy would be advantageous given the broad overlap of dangerous and benign-source disturbance amplitudes. *A. callidryas* combine amplitude with temporal and frequency information to refine their hatching decisions. The complexity of *A. callidryas* embryos’ risk assessment strategy is consistent with expectations based on the ambiguity and overlap in available cues, the high cost of both false alarms and missed cues, and the high mortality rates that create an opportunity for strong selection. It contrasts, however, with *a priori* expectations based on their developmental immaturity. Our growing understanding of embryo behavior in this and other species (Warkentin and Caldwell, 2009) suggests that, in the relevant ecological context, early life stages can evolve environmentally-cued responses as sophisticated as many we see in more mature organisms.

